# New records of Atlantic humpback dolphin in Guinea, Nigeria, Cameroon and Togo underscore fisheries pressure and generalized marine bushmeat demand

**DOI:** 10.1101/035337

**Authors:** Koen Van Waerebeek, Michael Uwagbae, Gabriel Segniagbeto, Idrissa L. Bamy, Isidore Ayissi

## Abstract

In northern Guinea, we sighted two groups of *Sousa teuszii* (n=25; n=40 dolphins) off the Tristao Islands during exploratory small-boat surveys in 2011–12. Based on these and recent (2013) observations in the contiguous Río Nuñez estuary, we propose a single ‘Guineas stock’, combining the former ‘Rio do Jêba-Bijagos’ and South Guinea stocks. Significant mortality of *S. teuszii* from fisheries interactions is widely recognised however not quantifiable as monitoring effort is sporadic. In Guinea, catches were documented in 2002 (n=1) and in 2011–12 (n=5). Landed specimens were recorded in Cameroon (n=2) and Nigeria (n=2). All individuals were killed in small-scale coastal fisheries, presumably as accidental net entanglements, though directed takes cannot be excluded. All landed dolphins were butchered for human consumption (marine bushmeat). Nigerian fishers indicated also an alternative use as shark bait. If local markets in cetacean bushmeat and bait develop, as in Ghana, that will exacerbate pressures by encouraging directed takes. Catch records in Nigeria and sightings in Togo authenticate both nations as (long-suspected) range states for *S. teuszii*, a belated documenting of the primary, historical distribution. The Gulf of Guinea stock (‘Cameroon dolphins’) extends at least from Togo to southern Cameroon, and probably into Equatorial Guinea. However, rare sightings of small groups point to remnant, not thriving, dolphin communities. We anticipate *de novo* distribution gaps emerging and consolidating, following decades of fisheries interactions and creeping encroachment on once pristine coastal habitat. Developed coastlines in Ghana and Côte d’Ivoire devoid of records may already constitute such gaps. As the lack or scarcity of records warn about formidable challenges to the long-term survival of *S. teuszii*, innovative, workable protection measures are needed, soonest. We recommend the implementation of several new border-straddling marine protected areas (cf. Saloum Delta-Niumi National Park Complex) which could bring forth a major conservation effect. Binational involvement bears obvious advantages, from sharing responsibilities and allowing for larger protected areas. Suggested dolphin sanctuary examples could include MPAs straddling borders between Cameroon/Equatorial Guinea and Guinea-Bissau/Guinea-Conakry.

## 1. Introduction

As applies to any wildlife species, the effective management and conservation of obligate coastal and estuarine marine mammals, such as humpback dolphins (*Sousa* spp.), require a comprehensive understanding of spatial overlap of its distribution and habitat with that of areas of intense anthropogenic pressures. Knowledge of the primary distribution of the Atlantic humpback dolphin *Sousa teuszii* (Kükenthal, 1892), a middle-sized (adults ca. 270 cm) African delphinid, remains incomplete and even new range states are still being added. Endemic to the subtropical and tropical eastern Atlantic, *S. teuszii* ranges exclusively in nearshore waters, apparently discontinuously from Dakhla Bay (N23°54’,W15°46’) southeast to Tombua, southern Angola (S15°47’,11°47’E). A comprehensive status review (Van Waerebeek et al. 2003, 2004) confirmed nine range states and provisionally identified eight management stocks based on the documented distribution record: Dakhla Bay (Western Sahara), Banc d’Arguin (Mauritania), Saloum-Niumi (Senegal, The Gambia), Canal do Gêba-Bijagos (Guinea-Bissau), South Guinea (Guinea), Cameroon Estuary (Cameroon), Gabon and Angola. Later field work confirmed continued presence in Cameroon after 110 yrs without a single record (Ayissi et al. 2011, 2014) and provided an improved understanding of occurrence in Gabon (Collins et al. 2004, 2010; Collins 2012, 2015) and Angola (Weir 2009; Weir et al. 2011). Recent records evidenced the occurrence of *S. teuszii* in the Republic of the Congo (Collins et al. 2010; Collins 2012), Benin (Zwart and Weir 2014), Nigeria (this paper) and Togo (this paper)^1^, bringing the count of confirmed range states to 13. Admittedly, several were long-suspected distribution areas. No records exist for Sierra Leone, Liberia, Côte d’Ivoire, Ghana and Equatorial Guinea. Considering neighbouring range states, *S. teuszii* is likely present in Equatorial Guinea (mainland Río Muni, not offshore Bioko), the Democratic Republic of the Congo (around the Congo River estuary) and Sierra Leone. Small-scale distributional information remains entirely fragmentary, and often dated, due to a persisting paucity of dedicated survey effort.

Provisionary stocks were defined for management purposes without evidence for distinct biological populations (Van Waerebeek et al. 2003, 2004). The historical distribution range of *S. teuszii* may remain uncertain considering that the comtemporary range may already have been modified by human factors. No robust abundance estimates are available for any stock, at best minima from added group sizes have been listed (reviewed in Collins 2015). Several stocks (e.g. Dakhla Bay, Banc d’Arguin, Angola) are extremely small, perhaps do not exceed a few tens of individuals (Van Waerebeek et al. 2003, 2004; Weir et al. 2011; Collins 2015) and hence risk extirpation.

Circumstantial evidence, including great distance (>350 km) and lack of sightings between core areas, suggests that the Dakhla Bay, Banc d’Arguin and possibly Siné-Saloum stocks may currently be reproductively isolated from each other and from more southern stocks (Van Waerebeek et al. 2004; Collins 2015). It is unclear whether the longest coastline without any *S. teuszii* records, stretching 1,900 km over four nations (Sierra Leone, 402 km; Liberia, 579 km; Côte d’Ivoire, 380 km; Ghana, 539 km) is an effective distribution gap or may be explainable by a lack of reporting. Lack of records for Ghana’s coast may be the result of extensive dolphin exploitation, and *S. teuszii* may now either be absent or extremely rare. If the latter, the Volta River delta (eastern Ghana) has been identified as a potential remnant habitat (Van Waerebeek et al. 2004, 2009; Van Waerebeek and Perrin 2010; Debrah et al. 2010). Based on credible fishermen’s reports, Van Waerebeek et al. (2003, 2004) suggested a probable Togo stock, however its existence remained unconfirmed for a decade (Segniagbeto et al. 2014) until finally photographed in 2014. In Benin, four individuals were sighted 14.5 km west of Cotonou port on 3 November 2013 (Zwart and Weir 2014), the only indication of occasional presence on Benin’s 121 km coast (see Sohou et al. 2013).

Captures in coastal fisheries and all types of coastal development are considered the principal threats to *S. teuszii* (Van Waerebeek 2003, 2004; Van Waerebeek and Perrin, 2007; Ayissi et al. 2011, 2013; Weir et al. 2011; Collins 2015). Several authors argued that the conservation status of *S. teuszii* seemed to have significantly deteriorated and that the IUCN Red List classification (“Vulnerable”, VU) should be modified (Van Waerebeek and Perrin 2007; Weir et al. 2011, Ayissi et al. 2013). Collins (2015) applying the Red List criteria suggested that the species merits a status classification of “Critically Endangered” (CR).

We here document recent records in four West African nations (Guinea, Togo, Nigeria, Cameroon) which underscore the common mortality in small-scale coastal fisheries and the high, possibly increasing, local demand for marine bushmeat derived from captured dolphins.

## 2. Material and Methods

Specimen and sightings data were obtained through either dedicated field effort in Nigeria and Guinea, or through opportunistic observations, in Togo and Cameroon.

In Nigeria, one of us (MU) surveyed Brass and Akassa Islands in Bayelsa State, Niger Delta region, between August 2011 and May 2012 in search of evidence of fisheries interactions with small cetaceans (and sea turtles), and in particular with *S. teuszii*. Field work was hampered due to the volatile nature of the region, including militant group activities. During courtesy visits, MU interviewed 48 artisanal fishermen and 2 fish mongers to learn their perspectives, evaluate their awareness of humpback dolphins and gather evidence on any cetacean catches. Several other fishermen were also informally queried. When interviewed, fishermen were shown colour photographs of Atlantic humpback dolphins, common bottlenose dolphins and humpback whales. Main focus areas included Rotel Fishing settlement (N04°19.45’, E06°14.79’) and Imbikiri Quarters, Twon Community (N04°17.54’, E06°17.41’). The 50 interviewees included about proportionally the active fishermen’s age classes, as follows: 21–30 yrs old (10%), 31–40 (54%), 41–50 (30%), >50 (6%).

Bamy (2011), guided by the extensive shallow water biotope in northern Guinea (Doumbouya and Camara, 2009), prime habitat for humpback dolphins, first confirmed *S. teuszii* near the Tristao Islands, a marine protected area at Guinea’s border with Guinea-Bissau. Two of us (ILB, KVW) undertook a second exploratory survey of the islands 4–9 June 2012 (Figure 1), nearshore from a 15hp outboard-powered canoe and through shore-based observations (Table 1). Visual search effort was both by naked eye and 7×50 binoculars. Canoe-based observer effort (mean v= 10.61 km h^−1^) totalled 974 min (16h,14min) and covered 172.28 km. Two fishermen acted as accessory observers. Shore-based surveys, on foot (675 min; 35.45 km), comprised visual scanning of the inter- and infratidal coastal strip up to 2 km from shore (Table 1). Flotsam accumulated at the high tide line was searched for cetacean remains. Fishermen at the Tristao Islands were queried about dolphin bycatches. Considering the area is a national Marine Protected Area (MPA) and non-residents require an access permit, the Tristao Islands enjoy a certain level of protection from disturbance.

**Figure 1.**
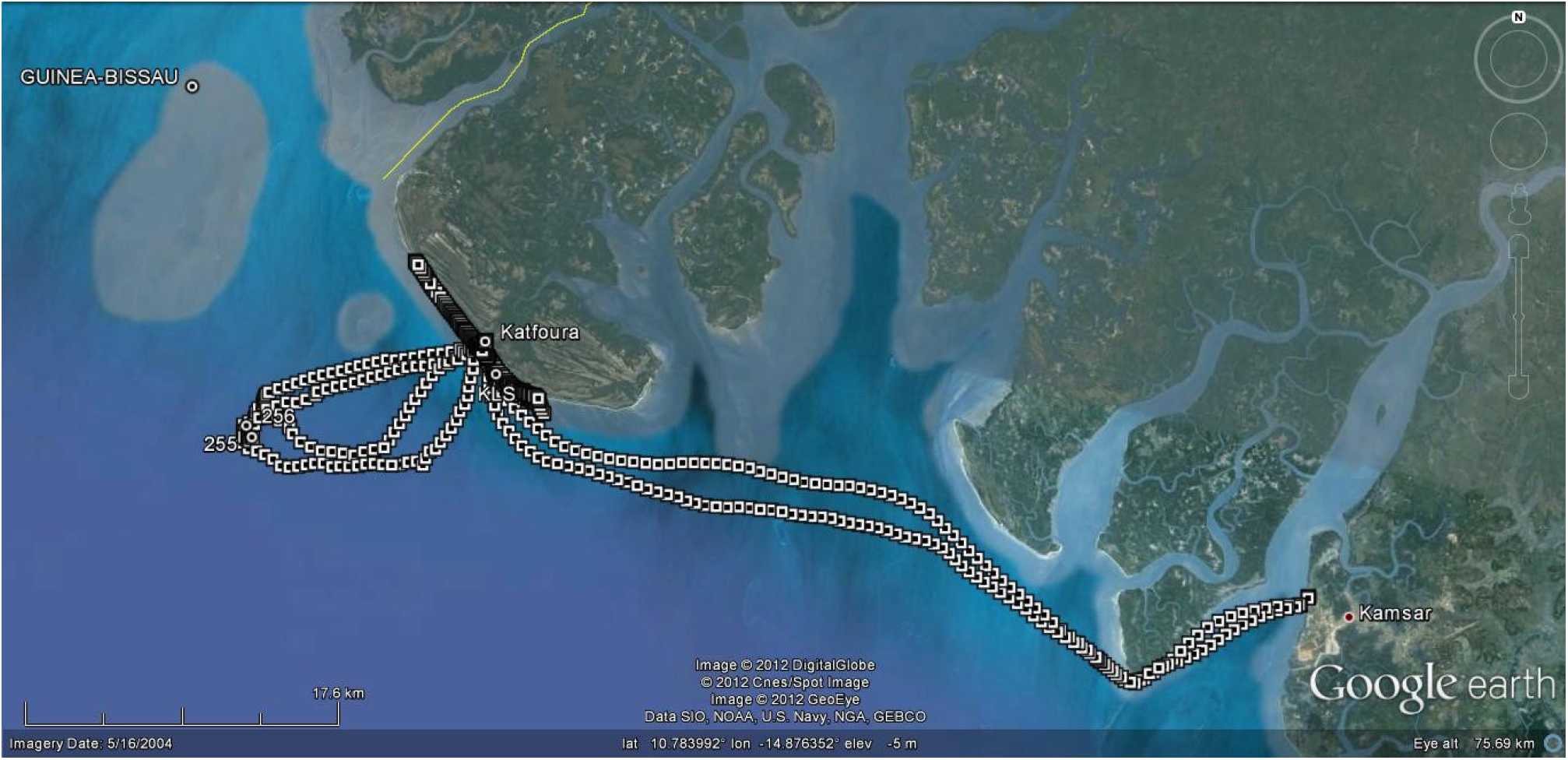
Map of the Tristao Islands study area, at the border between Guinea and Guinea-Bissau (yellow line). Indicated are survey trackline positions (white squares) between Kamsar and Katfoura and off Katfoura Island, and shore-based survey area (dark line). Sighting location of Atlantic humpback dolphin group (waypoint 255) is at the western extreme of tracklines. The dolphins moved in NW direction.

Members of the Togolese Society for Nature Conservation (Agbo-Zegué) including one of us (GS), conducted unstructured interviews in July 2013 and October 2014, querying coastal inhabitants about the presence of dolphins along Togo’s 50 km of mostly sandy beaches (Bight of Benin). Tourist operator Mr. Loïc Henry who, since 2010, organises year-round small-boat and fishing excursions in Togo’s coastal waters, was also interviewed.

In Cameroon, the Association Camerounaise de Biologie Marine (ACBM), led by IA, supports a small-scale informal network of observers, and opportunistically receives information about captures and strandings of sea turtles and cetaceans. No effort estimate can be linked to this all-volunteer programme.

**Table 1.** Coastal survey effort at Tristao Islands, Guinea, 5–10 June 2012. See Figure 1.

| Date | Survey effort | Comments |
| --- | --- | --- |
| 05 June | Small-boat survey Leg 1: Kamsar to Katfoura (via Nafaya) with pirogue. 08:20-12:45 (265min); odometer 52.08km. | No cetaceans sighted. |
| 06 June, AM | Katfoura beach survey, 07:40-11:00h; odometer 12.15km; beach-combing (on-foot) for specimens and nearshore sightings | No cetaceans sighted; no skeletal remains found. |
| 06 June, PM | Beach survey for nearshore sightings and cetacean remains: Katfoura direction SE, 12:00-17:00h, odometer 12.1km | Calvaria of juvenile *S. teuszii* recovered on beach between Katfoura and Nafaya |
| 07 June, AM | Small-boat survey Leg 2: depart Katfoura, 07:30-11:05h (215min), odometer 34.10km | Sighting #255 (re-sighting #256): 40 (30-45) *S. teuszii* |
| 07 June, PM | Visit of Katchek fishing port (Bamy): interviewing artisanal fishermen | Two skulls of *S. teuszii* (said to be 'by-catches') recovered from fishermen. |
| 08 June AM | Small-boat survey Leg 3: Katfoura direction south, in pirogue, 08:05-11:41h (216min), odometer 33.8km. | No cetaceans sighted. |
| 08 June PM | From Katfoura direction SE, 14:35-17:30h, odometer 11.2km (low tide). beach-combing (on foot). | No cetacean material found. |
| 09 June, AM | Small-boat survey Leg 4: Katfoura to Kamsar, 07:40-12:18h (278min); odometer 52.3km | No cetaceans sighted. |

## 3. Results and Discussion

### 3.1 Guinea

*Historical data*. Cadenat (1956) first observed and reported *S. teuszii* from Guinea, in inshore silt-laden waters south of Conakry, in January 1953, which formed the rationale for the definition of the ‘South Guinea’ management stock (Van Waerebeek 2003; Van Waerebeek et al. 2003, 2004).

*Recent sightings*. On 1 September 2011, Bamy (2011) first sighted a group of ca. 25 Atlantic humpback dolphins in shallow waters near the Tristao Islands, northern Guinea. On 7 June 2012 (at 09:15) two of the authors (KVW, ILB) observed a large, scattered group of ca. 40 (30–45) humpback dolphins at N10°45.332’, W15°08.912’ off Katfoura Island, one of the Tristao Islands (Figure 2A). At 09:40 (N10°45.677’, W015°09.080’) the group dispersed as the dolphins actively avoided the canoe, typical behaviour in *S. teuszii* (Van Waerebeek et al. 2003). Mean Lower Low Water depth in the area varied from 1–3 m, as read from a detailed Soviet chart (N°34406-G, published 15 December 1979, Département Principal de la Navigation et de l’Océanographie, Ministère de Défense de l’URSS). With high tide at 10:58 and a mean tidal range of 5 m, depth at both locations was estimated at 4.75–6.75 m. Turbidity was high. Multidirectional movements in subgroups of 2–4 individuals with frequent and abrupt speed changes pointed to active foraging during rising tide, observed also in Senegal’s Saloum Delta (Van Waerebeek et al. 2003, 2004). The great variability in shape and size of dorsal humps and fins, reflecting ontogenetic and sexual variation, indicated a mixed-age group with juveniles, adults (Figure 2B) and a single calf. In the largest individuals, the massive, quasi rectangular dorsal humps, raised about equally craniad as caudad, almost obfuscated the small dorsal fin proper. Posteriorly, slightly southeast of the Tristao Islands, five sightings were reported around the Río Nuñez Estuary (Weir 2015).

**Figure 2.**
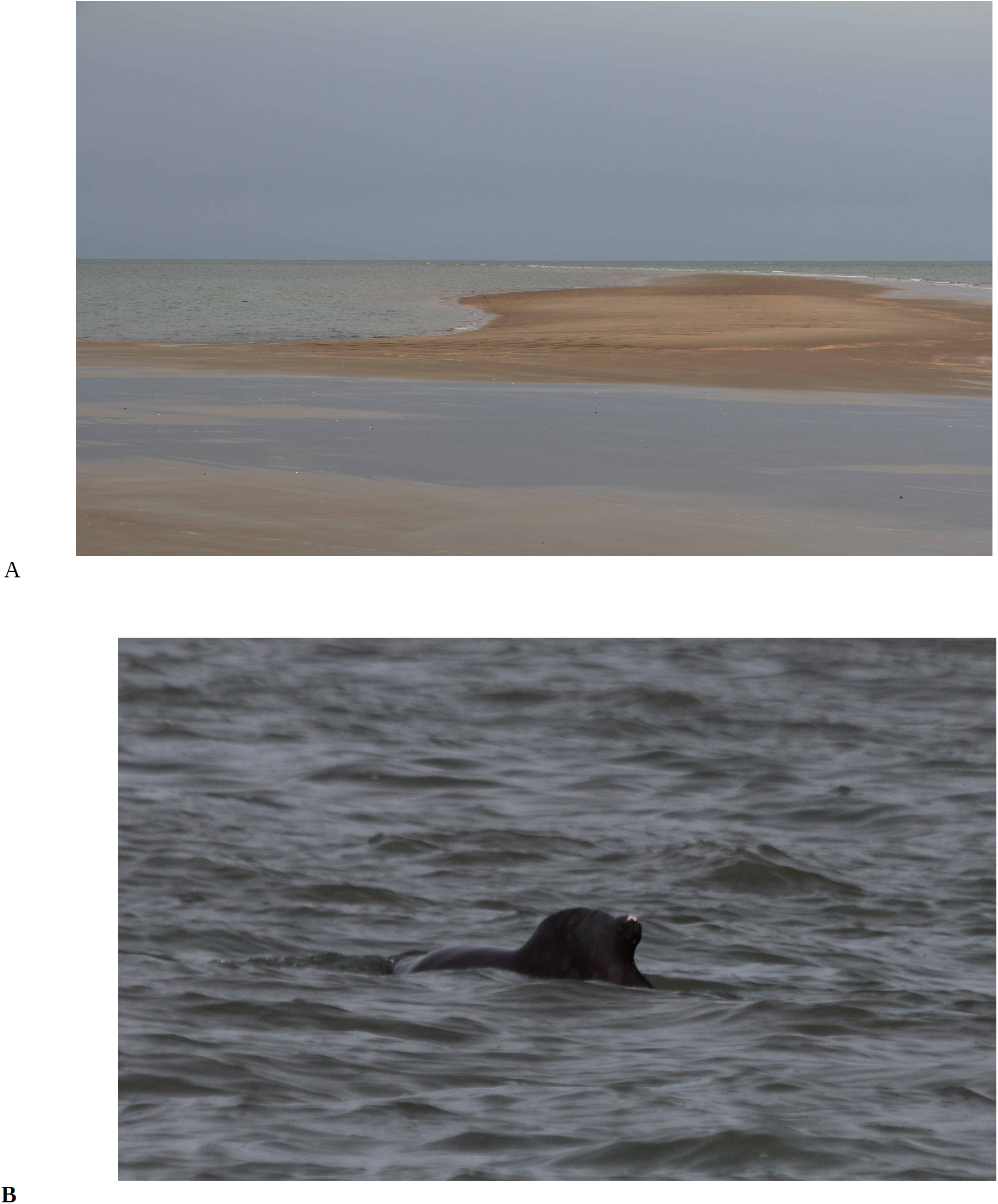
(A) Characteristic *S. teuszii* habitat at Katfoura Island, Tristao (here at low tide): shallow nearshore waters with strong tidal currents due to a maze of sand- and mudbanks and low-sloping sea bottom. (B) Earliest photographic record of live *S. teuszii* in Guinean waters. An adult member of a group of ca. 40 sighted at Katfoura Island, Tristao Islands, northern Guinea, on 7 June 2012. (Photos: K. Van Waerebeek).

*Fishery interactions*. The first documented *S. teuszii* specimen from Guinea consisted of a dolphin accidentally captured in an artisanal gillnet in the Bay of Sangaréah (Figure 3A), and landed at Dixinn on 13 March 2002 (Bamy et al. 2010; 2015). Bamy (2011) recovered cranial material of another two captured humpback dolphins at a fishing village on Tristao. On 6 June 2012, KVW collected a non-weathered calvaría (code 5 condition; Figure 3B) on the beach 1050 m east of Katfoura village (N10°48.215’,W015°01.754’) and 955 m west of Nafaya fishcamp (N10°47.692’, W015°01.424’). When shown at Katfoura, a fisher donated the matching mandibles and confirmed a recent capture. On 7 June 2012, ILB obtained the skulls of another two by-caught *S. teuszii* from fishermen at Kaatchek (N10°88.749’, W15°06.052’). These three dolphins, whose skulls are deposited at CNSHB, reportedly were consumed locally (marine bushmeat). We conclude that since 2002 there is material evidence for a minimum of six captures of *S. teuszii*.

**Figure 3.**
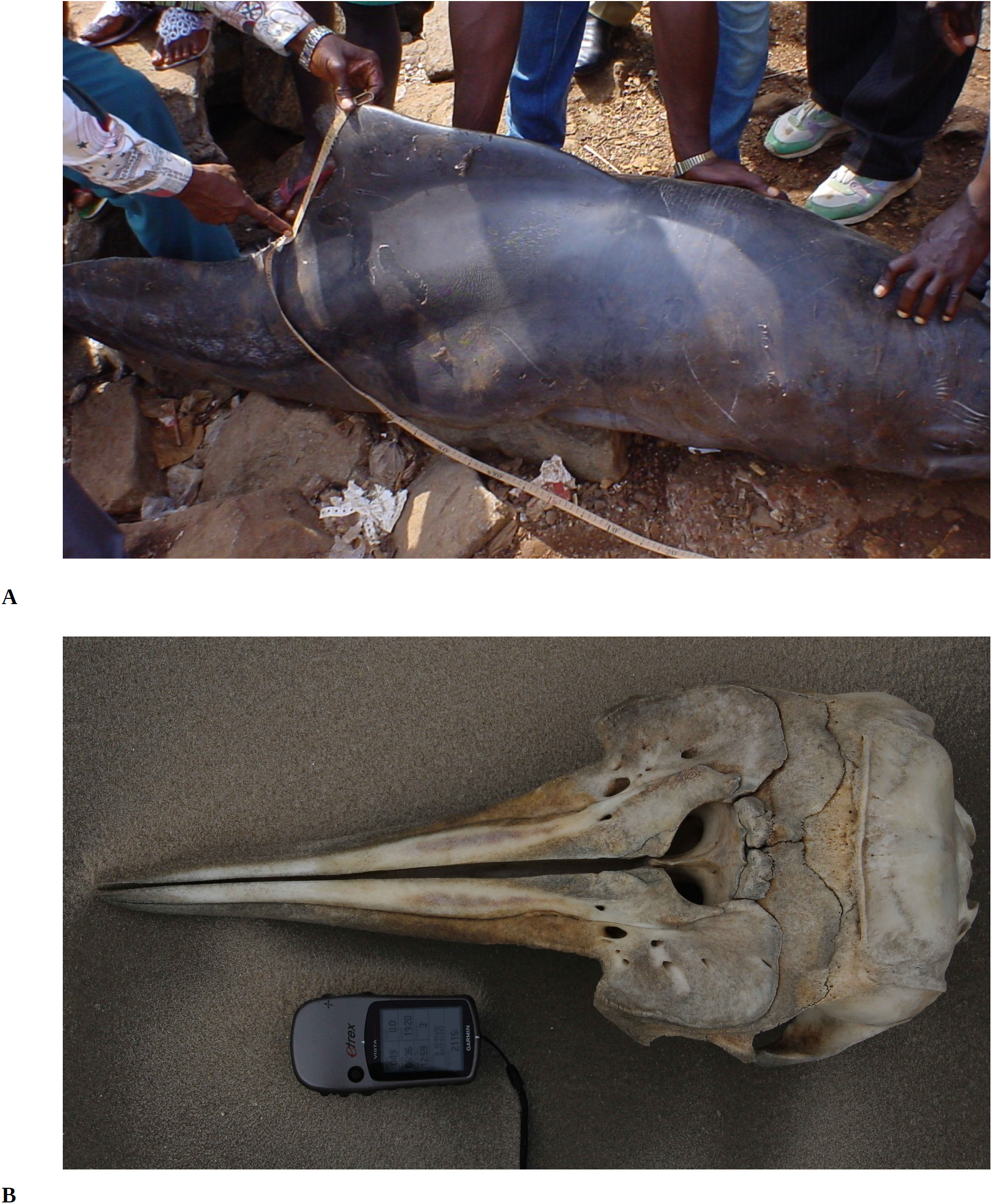
(A) Freshly captured, 222 cm male Atlantic humpback dolphin landed at the Dixinn fishing community, Guinea, on 13 March 2002. Note speckling on the tailstock and below the dorsal fin (Photo: I.L. Bamy). (B) Calvaría of juvenile *S. teuszii* collected between Katfoura and Nafaya fishing villages, at Katfoura Island, Tristao Islands, Guinea and a reported by-catch (see text). (Photo: K. Van Waerebeek).

*Folk knowledge*. Tristao fishermen referred to the specimens as “pueyegué” (Susu language) or also “furian” but it remained unclear whether these names refer to generic dolphins or specifically *S. teuszii*. The skull of a common bottlenose dolphin *Tursiops truncatus* was also collected at Tristao in 2011 (Bamy, 2011).

### 3.2 Togo

*Historical data*. Based on plausible, independent, reports from three senior fishermen interviewed at Lomé fish landing sites, Van Waerebeek (2003) first suggested the existence of a small population of Atlantic humpback dolphin (called ‘kposso’ by one fisher) off western Togo. This notion was reiterated later (Van Waerebeek et al. 2003, 2004; Segniagbeto and Van Waerebeek 2010; Segniagbeto et al. 2014). However, photographic evidence or scientific observations remained elusive for years. A first checklist of Togo’s cetaceans from authenticated stranding and bycatch records for 2003–2010 revealed nine cetacean species, but not *S. teuszii* (Segniagbeto and Van Waerebeek 2010; Segniagbeto et al. 2014).

*Sightings*. Mr. Loïc Henry (pers. comms to GS, March 2013 and *in litteris* to KVW on 02/04/2013 and 03/04/2013) photographed^2^ small groups of dolphins from shore (Figure 4) on four occasions (with estimated group sizes): on 27 December 2008 (n=7) at “some 30 km [NE] of Lomé” ca. N06.1953°, E01.4969; on 6 April 2013 (n=6) from a beach at Kpémé (N06.2253°, E01.5240°) on Togo’s centre coast; on 19 October 2013 (n=7) at Aného (ca. N06.2204°, E01.6141) and on 11 October 2014 (n=27, in several subgroups), locality unspecified. At the species identification request by Mr. Loïc Henry of these and other cetacean sightings, KVW (*in litteris* to LH, 03/04/2013) identified the dolphins discussed here as *S. teuszii* from the photographs provided. The sporadic occurrence of sightings (4 over a 7-year period) and small group sizes point however to low abundance. Also, till date, no specimen records are available from Togo (Segniagbeto and Van Waerebeek 2010; Segniagbeto *et al*. 2014), Ghana (Ofori-Danson et al. 2003; Debrah et al. 2010; Van Waerebeek *et al*. 2010, 2014) or Benin (Sohou *et al*. 2013).

**Figure 4.**
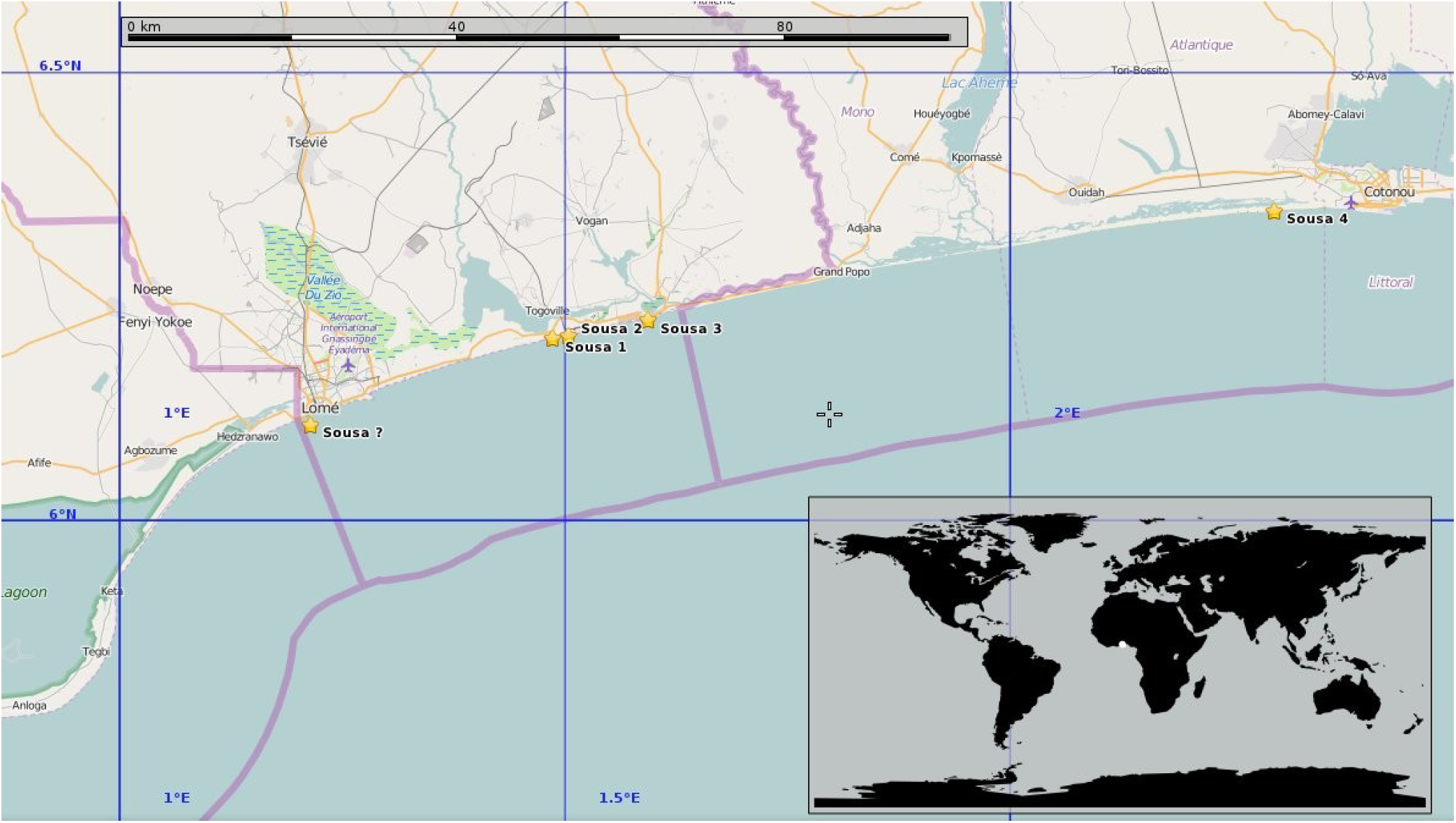
Map of northern Gulf of Guinea off Togo (centre), eastern Ghana (left) and western Benin (right). National borders indicated by pink lines. Shown are three confirmed sightings (Sousa 1–3) and one unconfirmed (Sousa?) sighting in Togo’s nearshore waters, and the only sighting (Sousa 4) reported for Benin (Zwart and Weir 2014). Animals are considered part of the Gulf of Guinea stock (Togo - Cameroon).

*Folk knowledge*. Fishermen reported inshore dolphins, likely *S. teuszii* and/or inshore *T. truncatus* from beaches around Benin’s Mono River estuary (Bouche du Roy) near Grand Popo, west to about Gbetsogbé (N06°09.135’, E01°18.333’), 5 km east of Lomé port. While traditional animist beliefs prevalent in Togo regard dolphins ‘totems’ to be protected from hunting or any harm (Segniagbeto *et al*. 2014), evidently these cannot prevent accidental net entanglements. In view of *S. teuszii*’s natural avoidance of human disturbances (Van Waerebeek et al. 2004; Weir et al. 2011), the lack of sighting records in the wider Lomé port area is thought to be linked to heavy vessel traffic, noise and chemical pollution as well as ongoing port construction work.

### 3.3 Nigeria

*Historical data*. Considering a 850 km coastline, much of it rimmed with creeks and dense mangrove forests, known habitat for *S. teuszii*, Nigeria has long been assumed a logical range state for the species (Klinowska 1991), critical reviews failed to identify any supported records (Van Waerebeek *et al*. 2004; Uwagbae and Van Waerebeek 2010; Collins 2015).

*Sightings*. The lack of authenticated sightings of *S. teuszii* in Nigeria, till date, can confidently be attributed to scarce scientific observer effort in coastal waters. Uwagbae and Van Waerebeek (2010), reviewing Nigerian cetaceans, found supporting evidence for only two cetacean species (*T. truncatus* and *Megaptera novaeangliae*) plus a credible but unsupported offshore record of *Lagenodelphis hosei*.

*Fisheries interactions*. Two dolphins killed in artisanal gillnets off Brass Island, Niger Delta, have become the first firm evidence for *S. teuszii* for Nigeria. An adult female was landed at the Rotel fishing settlement (N04°19.45’, E06°14.79’), Brass Island, in November 2011 (Figure 5B) and a second (juvenile) animal, also taken by local fishermen was landed at Imbikiri quarters (N04°17.54’, E06°17.41’), Twon Community, Brass Island, in February 2012 (Figure 5A). Both animals were butchered for marine bushmeat. Moore *et al*. (2010) and Solarin (2010) suggested that small cetaceans were rarely caught in fishing gear deployed by artisanal and industrial fishermen in Nigerian coastal waters. Uwagbae and Van Waerebeek (2010) disagreed and suggested that dolphins are likely captured with some frequency with takes going unreported, now corroborated by the Brass Island examples. Systematic, year-round monitoring will be required to gather statistics that reflect real mortality levels.

**Figure 5.**
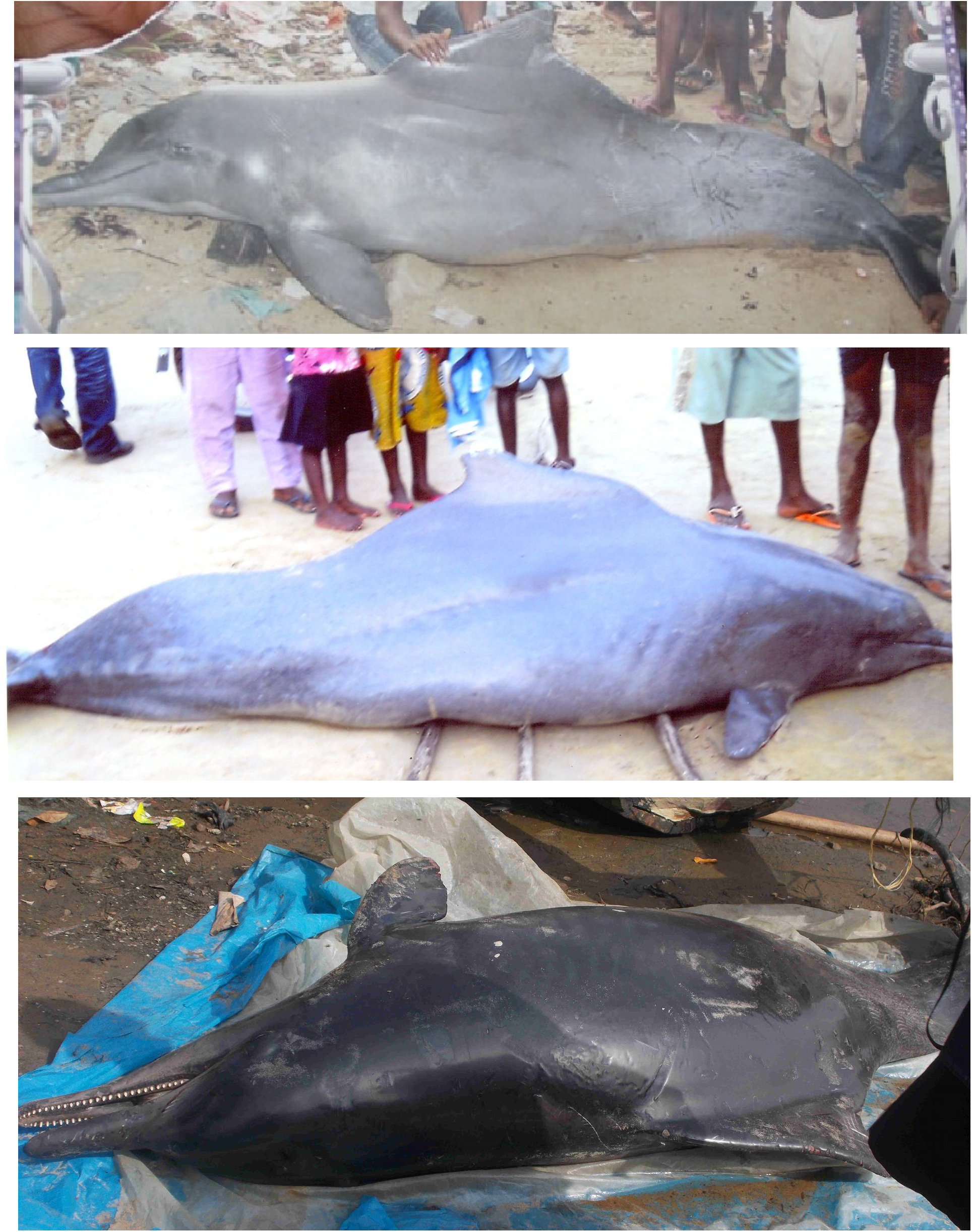
Atlantic humpback dolphins captured in artisanal coastal fisheries. (A) Juvenile landed at Imbikiri quarters, Brass Island, Nigeria, February 2012; (B) Adult specimen at Rotel fishing settlement, Nigeria, in November 2011. (Photos: Michael Uwagbae). (C) Freshly captured specimen landed at Londji landing site, Cameroon. (Photo: Isidore Ayissi).

*Interviews*. Among 50 interviewees on artisanal fishing practices, 66% indicated their professional activities included also ‘shark hunting’, 14% were also ‘shrimp fishing’ and 20% also engaged in subsistence farming. All respondents had sighted dolphins either at sea or from the beach. When at sea, 96% sighted dolphins ‘once in a while’ and 4% sighted them ‘very often’ (almost every fishing trip). Most fishermen (94%) admitted that at least once they had caught a dolphin, while 6% denied this. On the question whether they had encountered an accidentally entangled dolphin in their fishing gear, 58% replied ‘occasionally’ and 42% said ‘never’. When queried whether they had intentionally captured dolphins, 58% confirmed, 36% denied and 6% were ‘not sure’, meaning they tried but had given up when the hunt proved too difficult. Asked what they did with captured dolphins, 92% of respondents said it was used as bait to catch sharks, and 8% used or sold them for food. The retail value (in USD) of dolphins ranged widely, from 312–375 (22%), 440–500 (28%), 563–625 (16%) to >625 (34%). When asked when fishermen started hunting and trading in dolphins, they unanimously replied that it was an old tradition. However, this would imply that the utilisation of dolphins as marine bushmeat is far more prevalent than the interview data suggest, for the use as (shark) bait is a relatively recent usage and may have been an occasional event before shark fins were being exported to Asian markets. All fishermen recognised the Atlantic humpback dolphin and the common bottlenose dolphin from photographs shown, not surprisingly as both species are known to be taken (Uwagbae and Van Waerebeek 2010; this paper).

### 3.4 Cameroon

*Historical data*. The skull retrieved from a bycaught animal at Man O’War Bay, Cameroon, in 1892, curated at London’s Natural History Museum (BMNH 1893.8.11) (Jefferson and Van Waerebeek 2004), constitutes the holotype for *Sousa teuszii* (Kükenthal, 1892). Cameroon dolphin, a vernacular name used decades before the currently prevalent ‘Atlantic humpback dolphin’, is now applied solely to the Gulf of Guinea management stock (Van Waerebeek *et al*. 2004; Ayissi *et al*. 2014).

*Sightings*. On 17 May 2011, we observed a group of *ca*. 10 (8–12) Atlantic humpback dolphins near Bouandjo (N02°28.708’,E09°48.661’) in Cameroon’s South Region (Ayissi *et al*. 2014). The 259.1 km of visual small-boat survey effort in nearshore waters translated in an encounter rate of 0.039 individuals km^−1^. A 30.52 km shore-based survey yielded no sightings and no beach-cast specimen remains or strandings (Ayissi *et al*. 2014).

*Fisheries interactions*. A first capture of *S. teuszii*, supported by photographs, was landed by small-scale fishers at Campo, southern Cameroon, on an unspecified date in 2012 (Ayissi *et al*. 2014). An adult specimen landed at Londji fish landing site near Kribi (N02°56.1’, E09°54.6’) on 22 March 2014 (Figure 5C) reportedly became accidentally entangled in an artisanal gillnet in the Douala-Edea Fauna Reserve. No samples were collected as the dolphins were processed into marine bushmeat before researchers could reach the landing sites. These captures represent respectively the second and third confirmed *S. teuszii* specimens for Cameroon, after the holotype.

### 3.5 Population structure

The fact that large-ship platform-of-opportunity survey effort, covering 13,694 km in coastal waters of NW Africa in 2011–2013 (Djiba et al. 2015), yielded 270 primary sightings of 14 cetacean species but not a single humpback dolphin observation, strongly suggests that for all practical purposes *S. teuszii* is absent from shelf waters deeper than 20 m. This result is congruent with well-documented *S. teuszii* encounters, all of which occurred in shallow neritic waters (e.g. Van Waerebeek *et al*. 2004 and references therein; Weir et al. 2011; Ayissi et al. 2014; Collins 2015; this paper). Intraspecific morphological variation and molecular genetic population structure of *S. teuszii* has not been studied (Jefferson and Van Waerebeek 2004). Current distributional insights are incomplete due to a paucity of surveying, compounded by very low population numbers (e.g. Van Waerebeek 2003, Van Waerebeek *et al*. 2003, 2004; Weir et al. 2011; Collins 2015). If the absence of records between Sierra Leone and Togo would reflect an ecology-based distribution gap, then some population differentiation could be expected between NW African stocks and central African stocks. If however most of it is simply a very-low density zone, that recently emerged as a result of heavy anthropogenic pressures, then insufficient time would have passed for measurable population structure to have developed.

The new information presented herein calls for an update of some aspects of current insights of stock composition (Van Waerebeek *et al*. 2003, 2004; Van Waerebeek and Perrin 2007; Collins 2015). Van Waerebeek *et al*. (2004) provisionally defined two stocks in the Bissau/Conakry Guineas: the *South Guinea stock*, centred around Conakry, and the *Canal do Geba-Bijagos stock* of Bijagos Archipelago, Guinea-Bissau (Figure 6). The multiple specimen and sighting records from near the Tristao Islands, northern Guinea, in 2011–2012, as discussed above, as well as posterior sightings in the Río Nuñoz Estuary, SE of Tristao, in October-November 2013 (Weir 2015), are contiguous to the southern Guinea-Bissau border and evidently form part of the *Canal do Geba-Bijagos* stock. The Bay of Sangaréah, north of Conakry, is the northernmost documented location for the *South Guinea* stock (Van Waerebeek *et al*. 2004; Bamy *et al*. 2010). Since both areas are then separated by less than 150 km of relatively undeveloped coastline of mangroves and sandy beaches, it seems reasonable to assume that gene flow occurs, hence we here define a *Guineas stock* combining the former *Canal do Geba-Bijagos* and *South Guinea stocks* (Van Waerebeek *et al*. 2003, 2004). The humpback dolphins recorded in Togo, Benin and Nigeria are most likely “Cameroon dolphins”, *S. teuszii* communities belonging to the Gulf of Guinea stock, so defined based on the holotype and a 2011 sighting in Cameroon (Ayissi et al. 2014).

### 3.6 Fisheries impact

The consumption of bushmeat or wild meat is widespread in West Africa and embodies a complex and almost intractable problem once it acquires a (semi-)commercial scale (Brashares *et al*. 2004). The cultural and socio-economic drivers for the utilisation of cetaceans, manatees and sea turtles for human consumption bear many similarities to the terrestrial sourced bushmeat, which led to the introduction of the marine bushmeat concept (Alfaro-Shigueto and Van Waerebeek 2001; Clapham and Van Waerebeek 2007). In western Africa, the occasional or wide-spread consumption of cetacean bushmeat has been documented in an increasing number of countries, e.g. Senegal, The Gambia (Van Waerebeek et al. 2000, 2003, 2004; Murphy et al. 1997; Leeney et al. 2015), Guinea (Bamy et al. 2010, 2015; Bamy 2011; this paper), Ghana (Ofori-Danson et al. 2003; Debrah et al. 2010; Van Waerebeek et al. 2014), Togo (Segniagbeto and Van Waerebeek 2010; Segniagbeto et al. 2014), Benin (Sohou et al. 2013), Nigeria (Uwagbae and Van Waerebeek 2010; this paper), Cameroon (Ayissi et al. 2011, 2014), Gabon and Congo (Van Waerebeek and De Smet 1996; Collins 2012, 2015). An additional problem is the demand for dolphins to be used as bait in longline fisheries targeting sharks. This however appears relatively less prevalent than demand for marine bushmeat considering prices for the latter are high. In Ghana, for instance, cetacean bushmeat fetches the same prices as tuna species do.

All humpback dolphin specimens reported here, both freshly dead individuals and cranial material, were encountered in a context of artisanal fisheries, adding weight to the notion that local fisheries constitute the most acute threat to the Atlantic humpback dolphin (Van Waerebeek *et al*. 2004; Van Waerebeek and Perrin 2007; Weir *et al*. 2011; Ayissi et al. 2014; Collins 2015). When queried, locals typically admitted that dolphins were cut up and were either consumed or used as bait for shark fishing. It could not be determined whether landings of humpback dolphins originated primarily from directed or from incidental takes. Likely both occur in (at least) Ghana, Nigeria, Cameroon and Guinea.

Utilisation of *S. teuszii* as marine bushmeat is presently confirmed in four coastal nations, i.e. Cameroon, Nigeria (Ayissi et al. 2014; this paper), Guinea (Bamy et al. 2010; this paper), the Republic of the Congo (Collins 2015) and is suspected also for Senegal (Van Waerebeek et al. 2003) and possibly, in the past, Ghana (Van Waerebeek et al. 2004). However, no total mortality estimates from catches are available for any of the range states. Along parts of the Bight of Benin (Togo and Benin coasts), dolphins enjoy some protection against hunting thanks to indigenous animist beliefs among dominant ethnic groups, which may help explain why low numbers of *S. teuszii* are still present (this paper; Zwart and Weir 2014) and that no dead specimens have been found. However small-cetacean exploitation as prevalent in Ghana must warn against complacency. Many 100s of dolphins per annum of 14 different species are captured, openly landed and sold as marine bushmeat, albeit illegally, on an increasingly commercial scale (Van Waerebeek and Odori-Danson 1999; Ofori-Danson et al. 2003; Debrah et al. 2010; Van Waerebeek et al. 2014). While recognised in western Africa as an important challenge to marine mammal conservation (e.g. Ofori-Danson et al. 2013), fisheries authorities struggle to get a grip on the trade in dolphin products. In Ghana, up to now no working management programme is in place and all catch data have been collected and processed in an academic, not fisheries management, context. No Atlantic humpback dolphins have been recorded amongst multiple-year cetacean landings in Ghana, and no sightings were authenticated, which has been interpreted that either an ecological distribution gap exists (e.g. due to seasonal cool upwelling), or *S. teuszii* communities may have been depleted as a direct result of the dolphin exploitation and the coastal encroachment even before port monitoring effort started (Van Waerebeek *et al*. 2004, 2009; Weir *et al*. 2011). Some hope remains that rare observations of unidentified dolphins reported nearshore at the Volta River delta, eastern Ghana (Van Waerebeek et al. 2004) would prove to be *S. teuszii*. But then again, even if confirmed, chances are that it would comprise a remnant community, related or identical to those seen in Togo.

Although long-suspected, Nigeria and Togo have only recently been confirmed as range states, as were Congo (Collins et al. 2010) and Benin (Zwart and Weir 2014). Such findings, while evidently gratifying to the researchers involved, do not necessarily contain a reassuring conservation message. The scarce sightings in Benin and Togo rather carry a warning about a potential unsustainable situation. Overly optimistic claims^3^ that state or imply that an omnipresent *S. teuszii* occurs in a continuous, uninterrupted distribution are misleading and may compromise difficult conservation efforts that attempt to rally public awareness about an endangered marine mammal species. Extraordinary care must be taken in communications with the general public, with fisheries managers and decision makers lest misunderstandings grant licence to an already pervasive complacency in aquatic mammal management policies in many parts of Africa. Rare sightings of small groups of Atlantic humpback dolphins, e.g. in Benin (n=1 sighting; Zwart and Weir 2014), Togo (n= 4, this paper) and Cameroon (n=1, Ayissi et al. 2014) point to residual, not thriving, communities. We anticipate *de novo* distribution gaps emerging and consolidating, following decades of fisheries interactions and encroachment on once pristine coasts, reducing *S. teuszii’s* historical range. Irreversibly developed coastlines in Ghana and Côte d’Ivoire may already constitute *de novo* gaps, and the lack or scarcity of records warn about formidable conservation challenges. Since 2010, *S. teuszii* has been listed on both CMS Appendices I and II (Van Waerebeek and Perrin, 2007) and CITES Appendix I. Several authors (Ayissi et al. 2014; Weir et al. 2011) have indicated that the IUCN-allocated “Vulnerable” classification of *S. teuszii* is obsolete, as it does not reflect increased threat levels. Several of the management stocks are either already endangered or are heading into that direction. Collins (2015) proposed a “Critically Endangered” status which is meant to inform an upcoming review of the species’ IUCN Red Data Listing.

## Conclusions

‘New’ coastal range states are being named as researchers study recent field data from poorly studied shores, such as in the northern Gulf of Guinea (e.g. Zwart and Weir, 2014; this paper), essentially a belated documenting of the primary, historical range. While a practically-continuous distribution from Western Sahara to Angola, as often illustrated in cetacean guide books and suggested by Zwart and Weir (2015) may have existed along formerly pristine coasts, variably-sized *de novo* distribution gaps may be developing following decades of lethal fisheries interactions (Van Waerebeek et al. 2004; Van Waerebeek and Perrin 2007; Ayissi et al. 2014). The few surveys that provided density indicators have found very low abundance (reviewed in Collins 2015), i.e. few groups with small group sizes. The newly authenticated specimen records in Nigeria, Cameroon and Guinea underscore the significant mortality from bycatches and/or directed catches in local fisheries in parallel with a region-wide generalisation of demand for cetacean bushmeat, possibly related to diminished fish catches. Prey competition from fishers, habitat loss and related disturbance due to unstoppable coastal development (Van Waerebeek *et al*. 2004; Van Waerebeek and Perrin 2007; Weir *et al*. 2011; Collins 2015) jointly pose formidable challenges to the long-term survival of *S. teuszii*. Practicable protection measures are needed soonest. Several new border-straddling marine protected areas, similar to the Saloum-Niumi Complex uniting Senegal’s Saloum Delta National Park with The Gambia’s Niumi National Park (Van Waerebeek 2003), could have a major conservation effect as they can impede run-away coastal development. Bi-national involvement has obvious advantages in sharing tasks and responsibilities and allowing for larger protected areas. Also, border-straddling MPA’s may be more efficient since effective control of human transiting, limiting impact, is a key interest that coincides with those of authorities charged with guarding national borders. Proposals may include binational dolphin sanctuaries between Cameroon/Equatorial Guinea and Guinea-Bissau/Guinea-Conakry.

## Acknowledgements

Several people contributed to the success of the survey in Guinea, in particular we owe thanks to Mr. Abdoul Salam Bah, Conservator of the Tristao Islands MPA and to Mr. Mamadou Kaly Bah, former director-general of CNSHB for providing the ‘Ordre de mission’. We thank also the town elders and inhabitants of Katfoura. M. Uwagbae thanks Duke University Marine Laboratory and Oak Foundation (USA) for a study grant to cover field work in Nigeria. The efforts of community members of Brass and Akassa Islands and members of the Akassa Sea Turtle Foundation who provided information and support, are gratefully acknowledged. While in Guinea, KVW was a consultant for Intergovernmental Oceanographic Commission (IOC)-UNESCO. Boat rent and fuel in Guinea was kindly supported by a donation from Dr. Anna Vecchione, in honour and memory of late Mr. Edgar Quintel. CEPEC provided field equipment.

## Footnotes

1 Collins (2015) merely mentioned Togo for completeness but did not document sightings.

2 Photos are posted by LH (with public access) at https://www.facebook.com/media/set/?set=a.609858899141331.1073741829.585686184891936&type=3

3 e.g. Title “Filling in the gaps [in distribution]… ” (Zwart and Weir 2014) gives the misleading impression that *S. teuszii* occurs ubiquitously along all western African coasts.

